# Functional systemic CD4 immunity is a differential factor for clinical responses to PD-L1/PD-1 blockade therapy in lung cancer

**DOI:** 10.1101/508739

**Authors:** Miren Zuazo, Hugo Arasanz, Gonzalo Fernández-Hinojal, Maria Jesus García-Granda, María Gato, Ana Bocanegra, Maite Martínez, Berta Hernández, Lucía Teijeira, Idoia Morilla, Maria Jose Lecumberri, Angela Fernández de Lascoiti, Ruth Vera, Grazyna Kochan, David Escors

## Abstract

The majority of lung cancer patients progressing from conventional therapies are refractory to PD-L1/PD-1 blockade monotherapy. Here we show that baseline systemic CD4 immunity is a differential factor for clinical responses. Patients with functional systemic CD4 T cells included all objective responders and could be identified before the start of therapy by having a high proportion of memory CD4 T cells. In these patients CD4 T cells possessed significant proliferative capacities, co-expressed low levels of PD-1/LAG-3 and were responsive to PD-1 blockade *ex vivo* and *in vivo*. In contrast, patients with dysfunctional systemic CD4 immunity did not respond even though they had circulating lung cancer antigen-specific CD4 and CD8 T cells. CD4 T cells in these patients proliferated very poorly, strongly co-upregulated PD-1/LAG-3 after stimulation, and were largely refractory to PD-1 monoblockade. CD8 immunity only recovered in patients with functional CD4 immunity. T cell dysfunctionality was caused by PD-1/LAG-3 coexpression, and could be reverted by co-blockade. Patients with functional CD4 immunity and PD-L1 tumor positivity exhibited response rates of 70%, highlighting the contribution of CD4 immunity for efficacious PD-L1/PD-1 blockade therapy.

## INTRODUCTION

PD-L1/PD-1 blockade is demonstrating remarkable clinical outcomes since its first clinical application in human therapy (1,2). These therapies interfere with immunosuppressive PD-L1/PD-1 interactions by systemic administration of blocking antibodies. PD-L1 is overexpressed by many tumor types and generally correlates with progression and resistance to pro-apoptotic stimuli (3–5). PD-1 is expressed in antigen-experienced T cells and interferes with T cell activation when engaged with PD-L1 (6,7). The majority of advanced non-small cell lung cancer (NSCLC) patients progressing from conventional cytotoxic therapies who receive PD-L1/PD-1 blockade therapy do not respond. The causes for these distinct clinical outcomes are a subject for intense research (8). Emerging studies indicate that PD-L1/PD-1 blockade therapy does not only affect the tumor microenvironment, but also alters the systemic dynamics of immune cell populations (9–12). Some of these changes do correlate with responses and could be used for real-time monitoring of therapeutic efficacy. For example, PD-1^+^ CD8 T cells expand systemically after PD-1 blockade therapy in lung cancer patients (9). As CD8 T cells are the main direct effectors of responses through cytotoxicity over cancer cells, these changes are thought to be the consequence of efficacious anti-tumor immunity. Indeed, CD8 T cell infiltration of tumors correlates with good outcomes (13). However, the role of CD4 immunity in patients undergoing PD-L1/PD-1 blockade therapy remains poorly understood although extensive pre-clinical data link CD4 responses to anti-tumor immunity. Hence, CD4 T cells recognizing tumor neoepitopes contribute significantly to the efficacy of several types of immunotherapies in murine models and in cancer patients (14–16).

Human T cells undergo a natural differentiation process following the initial antigen recognition, characterized by the progressive loss of CD27 and CD28 surface expression, and acquisition of memory and effector functions (17,18). Hence, human T cells can be classified according to their CD27/CD28 expression profiles into poorly-differentiated (CD27^+^ CD28^low/negative^), intermediately-differentiated (CD27^negative^ CD28^+^) and highly-differentiated (CD27^negative^ CD28^low/negative^, T_HD_) subsets (17). Highly-differentiated T cells in humans are composed of memory, effector and senescent T cells, all of which could modulate anti-cancer immunity in patients and alter susceptibility to immune checkpoint inhibitors. To understand the impact of systemic CD4 and CD8 T cell immunity before the start of immunotherapies, we carried out a discovery study in a cohort sample of 51 NSCLC patients undergoing PD-1/PD-L1 immune checkpoint blockade therapy after progression to platinum-based chemotherapy. Our results indicate that baseline functional systemic CD4 immunity is required for objective clinical responses to PD-L1/PD-1 blockade therapies, with PD-1/LAG-3 co-expression causing CD4 and CD8 dysfunctionality in non-responder patients.

## RESULTS

### The baseline percentage of systemic CD4 T_HD_ cells within CD4 cells separates NSCLC patients in two groups with distinct clinical outcomes

To test the contribution of systemic T cell immunity to anti-PD-L1/PD-1 immunotherapy in NSCLC patients, a prospective study was carried out with a cohort of 51 patients treated with PD-L1/PD-1 inhibitors following their current indications (Supplementary table 1). These patients had progressed to conventional cytotoxic therapies. 78.4% presented an ECOG of 0-1, 70.6% with at least three affected organs and 25.5% with liver metastases (Supplementary table 1).

As a first approach, the relative proportion of CD4 T cell differentiation subsets (according to CD27/CD28 expression) within CD4 cells was quantified from fresh peripheral blood samples and compared to healthy age-matched donors. We could not analyze samples retrospectively as freezing and storage significantly altered surface expression markers in T cells.

Cancer patients showed a distribution significantly different from that of healthy donors in the percentage of CD4 T_HD_ cells (P<0.001) (Fig. 1A). Patients segregated in two groups separated by an approximate cut-off value of 40% CD4 T_HD_ (Fig. 1A); Patients with high (G1 group, 63.25±13.5 %, N=23) and with low percentages of baseline T_HD_ cells (G2 group, 27.05±10.6 %, N=28). Differences between G1 and G2 were also highly significant. Objective responders in our cohort study were found within G1 patients (P=0.0001) but not in G2 (Fig. 1A). In patients with measurable disease, significant regression of tumor lesions was only apparent for G1 profiles and progression for G2 patients (Fig. 1B). ROC analysis confirmed the highly significant association of baseline CD4 T_HD_ cells with clinical output (P=0.0003), providing a cut-off value of >40% to identify objective responders with 100% specificity and 70% sensitivity (Fig. 1C). A validation dataset from 32 patients was performed by parallel independent double-blind sample handling, staining, data collection and analyses (Supplementary Fig. 1). To remove background from CD4^low^ cells in the validation set, CD14^+^ monocytic cells were depleted before staining or gated out during analyses of flow cytometry data. Cohen’s kappa coefficient confirmed the agreement between the discovery and the validation sets of data on G1/G2 profile classification (κ=0.932). The highly significant association between G1 profiles and objective responses in the validation set was confirmed (P=0.0006) (Supplementary Fig. 1).

Progression-free survival (PFS) of patients stratified only by their baseline CD4 T cell profiles confirmed the association between G1 profiles and objective responses. The median PFS (mPFS) of G2 patients was only 6.1 weeks (95% C.I., 5.7 to 6.6) compared to 23.7 weeks for G1 patients (95% C.I., 0-51.7; P=0.001) (Fig. 1D). A comparison of G2 *versus* G1 baseline profiles showed hazard ratios for disease progression or death that favored the latter [3.1 (1.5-6.4; 95% C.I.) P = 0.002].

**Figure 1.**
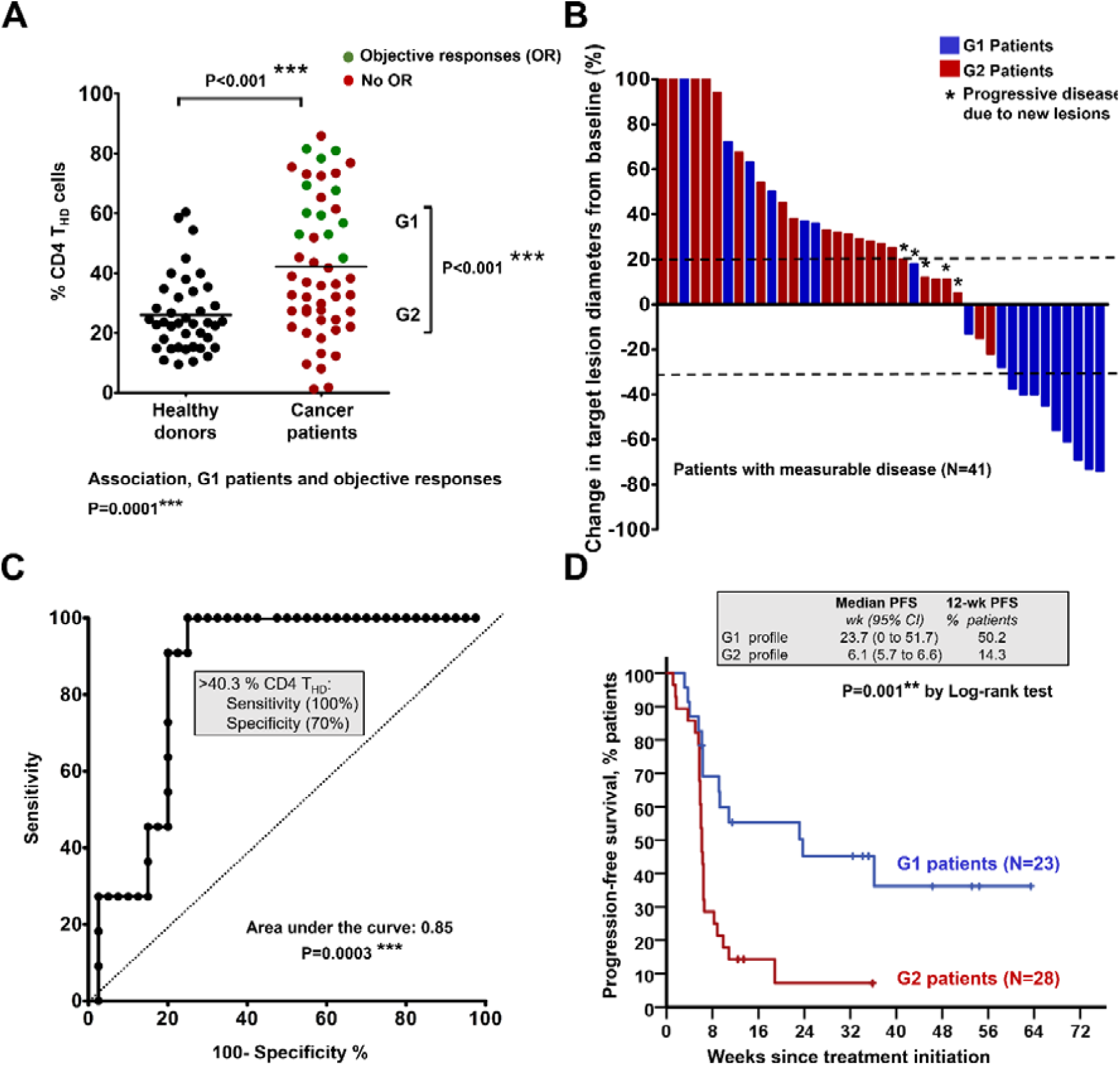
Baseline profiling of CD4 T cell differentiation subsets stratifies clinical responses to PD-L1/PD-1 blockade. **(A)** Percentage of circulating highly differentiated CD4 T cells within CD4 cells (upper graph) in age-matched healthy donors or NSCLC patients before undergoing immunotherapies. G1 and G2, groups of patients classified according to high T_HD_ cells (G1, >40% CD4 T_HD_ cells) and low T_HD_ cells (G2, <40% CD4 T_HD_ cells). Relevant statistical comparisons are shown by the U of Mann-Whitney test. In green, objective responders (OR). In red, no OR. Below the graph, correlation of objective responses to G1 and G2 groups by the Fisher’s exact test. **(B)** Waterfall plot of change in lesion size in patients with measurable disease classified as having a G1 (blue) or G2 (red) profile. Dotted lines represents the limit to define significant progression (upper line) or significant regression (lower line). **(C)** ROC analysis of baseline CD4 THD quantification as a
function of objective clinical responses. **(D)** Kaplan-Meier plot for PFS in patients treated with immunotherapies stratified only by G1 (green) and G2 (red) CD4 T cell profiles. Patients starting therapy with a G2 profile had an overall response rate (ORR) of 0% and 82% of them experienced progression or death by week 9. ORR was 44.8% for G1 patients, and the 12-week PFS was 50.2%.

To assess whether CD4 T cell profiling had prognostic value, overall survival (OS) of G1 progressor patients was compared to G2 patients. Compared to G2 patients, G1 progressors did not show survival advantage during immunotherapy (Supplementary Fig. 2). This was further supported by lack of significant association between responses and baseline ECOG score (P=0.6), although only a minority of our patients had ECOG >2. In our cohort there was no significant association between CD4 T cell profiles/objective responses with presence of liver metastases (P=0.88), or with tumor load (P=0.19). In agreement with all these results, we found no significant association of CD4 T cell profiles or objective responses with the Gustave-Roussy immune score (GRIm) (**P=0.14,** supplementary table 2), in patients with available data on number of metastases, serum lactate dehydrogenase (LDH) and neutrophil-to-lymphocyte (NLR) ratio (19). The hazard ratio for progression or death of G2 patients maintained its statistical significance by multivariate analyses (HR 9.739; 95% CI 2.501 to 37.929) when adjusted for tumor histology, age, gender, smoking habit, liver metastases, number of organs affected, PD-L1 tumor expression, NLR, serum LDH and albumin. Hence, we concluded that CD4 T_HD_ profiling has no significant prognostic value.

### Functionality of systemic CD4 immunity marks clinical outcomes and susceptibility to PD-L1/PD-1 blockade

Our data suggested that the separation of patients in two groups according to CD4 T_HD_ quantification reflected differences in the functionality of baseline systemic CD4 immunity before starting immunotherapy. First, we wondered whether PD-1 expression in non-stimulated circulating CD4 T cells was a differential factor between G1 and G2 groups. However, no differences were observed even when compared with healthy age-matched donors (not shown). We then tested if PD-1 was differentially upregulated after *ex vivo* stimulation with lung cancer cells. To this end and to standardize comparisons between donors, we engineered a human lung adenocarcinoma cell line (A549) to express a membrane-bound anti-CD3 single-chain antibody (A549-SC3 cells). This cell line stimulates T cells while preserving other cancer cell-T cell interactions such as PD-L1/PD-1 or MHC II-LAG-3 (supplementary figure 3A and 3B). This ensured the same standard assay for cancer cell-T cell recognition for each patient, with the same affinity and specificity (supplementary figure 3B, 3C and 3D).

CD4 T cells from NSCLC patients significantly up-regulated PD-1 compared to CD4 T cells from age-matched healthy donors after incubation with A549-SC3 cells (P<0.001) (Supplementary figure 3C and Fig. 2A). However, no significant differences in PD-1 expression were found between G1 and G2 CD4 T cells. As co-expression of PD-1 with LAG-3 is a trademark of dysfunctional human tumor-infiltrating lymphocytes in NSCLC (20), we studied PD-1 and LAG-3 co-upregulation in CD4 T cells from G1 or G2 patients. Accordingly, CD4 T cells from G2 donors presented a significantly higher percentage of T cells co-expressing both markers than G1 counterparts (Fig. 2B). To test whether G1 and G2 CD4 T cells showed differential proliferative capacities, the percentage of proliferating ki67+ cells was also assessed in a sample of patients (Fig. 2C). Accordingly, CD4 T cells from G2 patients were remarkably impaired in proliferation compared to their G1 counterparts after *ex vivo* activation with A549-SC3 cells. As G1 patients showed a higher percentage of baseline circulating CD4 T_HD_ cells compared to G2, we also tested whether this subset was responsive to activation by A549-SC3 cells (Fig. 2D). Interestingly, CD4 T_HD_ cells strongly proliferated in all patients although they constituted a minority in G2 patients. Hence, the strong proliferative capacities of CD4 T_HD_ cells in our study suggested that these were not senescent T cells, but highly differentiated memory subsets. To test this, their phenotype according to CD62L/CD45RA surface expression was assessed in a sample of patients before undergoing immunotherapies (Supplementary Fig. 4A). Compared to healthy donors, the majority of CD4 T_HD_ cells were comprised of central-memory (CD45RA^negative^ CD62L^+^) and effector-memory (CD45RA^negative^ CD62L^negative^) cells, without significant differences between G1 and G2 patients. Increased genotoxic damage is frequently associated to senescent T cells and can be used as a marker of senescence by evaluating the expression of H2AX (18). Interestingly, unlike CD4 T cell subsets from age-matched healthy donors, NSCLC CD4 T cells exhibited extensive genotoxic damage in both CD28^+^ and CD28^negative^ subsets without differences between G1 and G2 groups (Supplementary Fig. 4B). Therefore, genotoxic damage did not identify senescent T cells in patients progressing from conventional cytotoxic therapies. Then, the expression of the classical replicative senescence marker CD57 was used to identify senescent T cells, which accounted to about 30% of total T_HD_ cells in healthy age-matched donors, and about 10% in NSCLC patients (Supplementary Fig. 4C). Our results strongly suggested that circulating CD4 T_HD_ cells in our cohort of NSCLC patients mostly corresponded to non-senescent memory subsets.

**Figure 2.**
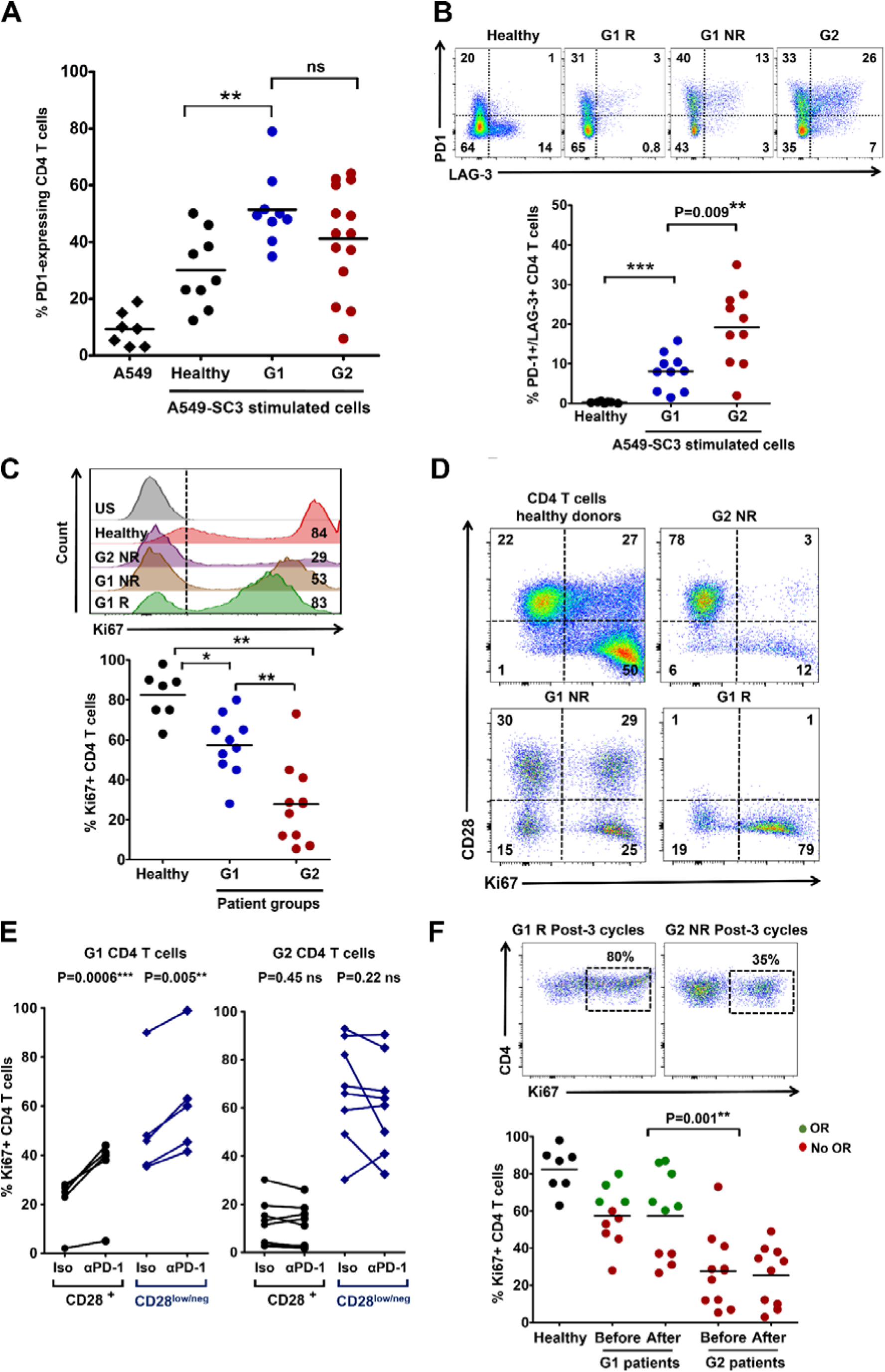
Differential systemic CD4 immunity and responses to PD-1/PD-L1 blockade in NSCLC patients. **(A)** The scatter plot shows PD-1 expression after co-culture of CD4 T cells from healthy donors or NSCLC patients, as indicated, with A459-SC3 lung cancer cells. Relevant statistical comparisons are indicated. **(B)** Upper graphs, flow cytometry density plots of PD-1 and LAG-3 co-expression in CD4 T cells from healthy donors, a G1 responder (G1 R), a G1 non-responder (G1 NR) and a G2 non-responder as indicated, following stimulation with A549-SC3 cells. Percentage of expressing cells are indicated within each quadrant. Below, same as in the upper graphs but as a scatter plot of the percentage of CD4 T cells that simultaneously co-express PD-1 and LAG-3. Relevant statistical comparisons are shown. **(C)** Upper flow cytometry histograms of Ki67 expression in CD4 T cells from the representative subjects as indicated on the right, after stimulation with A549-SC3 cells. Vertical dotted line indicates the cut-off value of positive vs negative Ki67 expression. The percentage of Ki67-expressing CD4 T cells is shown within the histograms. Below, same data represented as a scatter plot from a sample of G1 and G2 donors as indicated, with relevant statistical comparisons. **(D)** Proliferation of CD4 T cells stimulated by A549-SC3 cells from the indicated patient groups. CD28 expression is shown together with the proliferation marker ki67. Percentages of cells within each quadrant are shown. **(E)** Same as in (D) but in the presence of an isotype control antibody or an anti-PD-1 antibody with the equivalent sequence to pembrolizumab. The effects on CD4 T cells from a G1 and a G2 patient are shown, divided in CD28 high or low/negative subsets as indicated. Relevant statistical comparisons are shown. **(F)** Top, flow cytometry density plots of Ki67 expression in CD4 T cells from representative G1 or G2 patients after three cycles of therapy, activated by incubation with A549-SC3 cells. Below, same as above but as a dot-plot graph. A comparison between proliferating CD4 T cells before and after therapy is shown in unpaired patient samples. G1 R, G1 objective responder patient. G1 NR, G1 patient with no objective responses; Green, objective responders (OR) and red, no OR; Iso, treatment with an isotype antibody control; α-PD-1, treatment with anti-PD-1 antibody;*, **, *** represents significant (P<0.05), very significant (P<0.001) and highly significant differences (P<0.0001), respectively.

Considering the distinct patterns of PD-1/LAG-3 expression between G1 and G2 CD4 T cells, we tested whether they responded differentially to *ex vivo* PD-1 monoblockade. Hence, CD4 T cells from a sample of G1 and G2 patients were co-cultured with A549-SC3 in the presence of an anti-PD1 antibody molecularly equivalent to pembrolizumab (21) (Fig. 2E). PD-1 blockade increased proliferation of G1 CD28^+^ and CD28^negative^ CD4 T cell subsets, while their G2 counterparts were largely refractory. To find out if G2 CD4 T cells were unresponsive to PD-1 blockade *in vivo*, CD4 T cells from G1 and G2 patients were obtained after at least three cycles of therapy and tested for their proliferative capacities (Fig. 2F). Proliferative differences between G1 and G2 CD4 T cells were still maintained during immunotherapy.

We wondered whether G2 patients had also decreased CD4 pre-immunity to lung cancer cells before the start of immunotherapy that could contribute to the lack of responses in this group. CD4 T cells specific for lung adenocarcinoma-associated antigens were quantified in peripheral blood using a presentation assay with autologous monocyte-derived DCs as described (22). DCs were loaded with A549 cell lysate as a source of common human lung adenocarcinoma antigens (23), as in most instances we did not have sufficient biopsy material as a source of autologous tumor-associated antigens (TAAs), or to quantify/isolate tumor-infiltrating CD4 cells. TAA-loaded and control unloaded DCs were matured with IFN-γ to make sure that even strongly dysfunctional T cells could be activated. Lung cancer cell-reactive CD4 T cells were identified by co-expression of IL-2 and IFN-γ (Fig. 3A). Interestingly, TAA-specific CD4 T cells could be detected in circulating CD4 T cells in G1 and G2 patients before the start of immunotherapy, even at high proportions in some patients. Indeed, the relative percentage of circulating TAA-specific CD4 T cells was not significantly different between a sample of G1 (responders and non-responders) and G2 patients (Fig. 3B), at least using A549 lung cancer cell antigens as comparative standard. To find out whether these TAA-specific CD4 T cells were preferentially found within T_HD_ or non-T_HD_ populations, CD28 expression was evaluated. Interestingly, TAA-specific CD4 T cells consisted of both T_HD_ and non-T_HD_ subsets, without significant differences in relative percentages between G1 and G2 patients (Fig. 3B). These results suggested that poor responses in G2 patients in our cohort study were not caused by absence of tumor-specific CD4 T cells but rather by intrinsic dysfunctionality.

**Figure 3.**
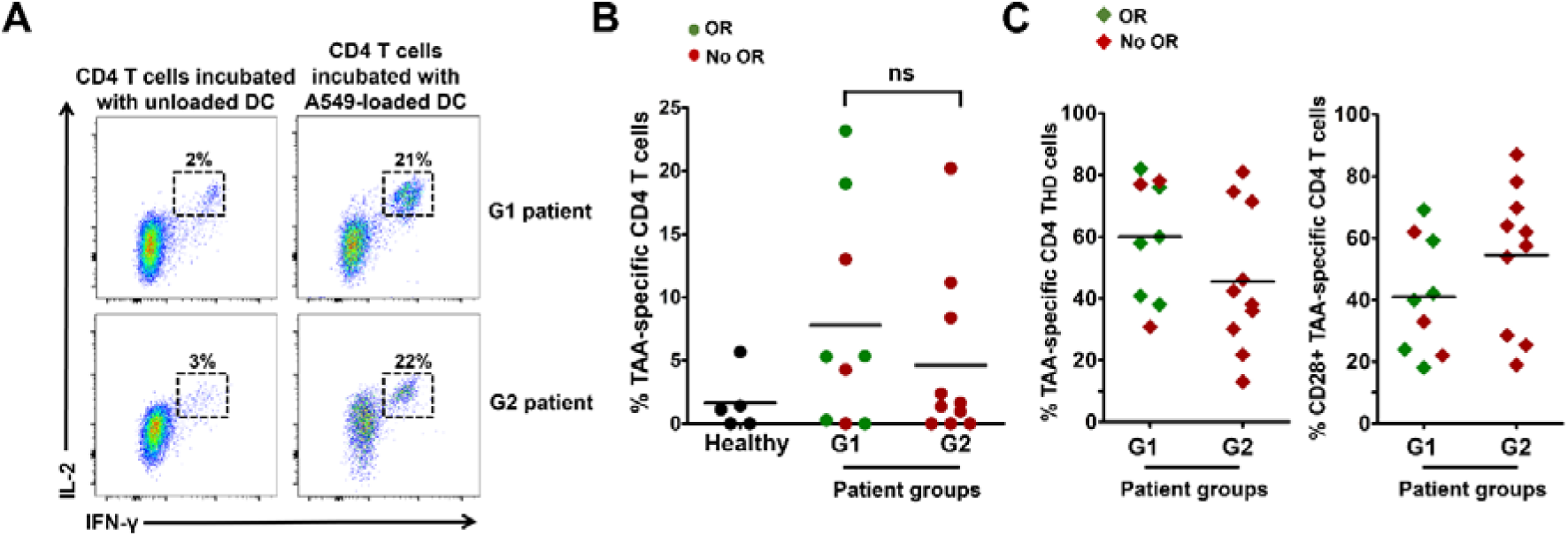
Lung cancer antigen-specific CD4 T cells in NSCLC patients. **(A)** Expression of IFN-γ and IL-2 in CD4 T cells from G1 or G2 representative patients incubated with A549 antigen-loaded autologous DCs or with unloaded controls activated with IFN-γ as indicated. Percentages of events are shown within each quadrant. **(B)** as in (A) but representing the data from a sample of G1 and G2 patients as a scatter plot graph. Objective responses (OR) are shown in green. In red, patients with no OR. **(C)** The scatter plot graph on the left represents the percentage of CD4 T_HD_ cells within lung-cancer specific CD4 T cells in a sample of patients from the indicated G1/G2 groups. On the right, same as left but representing the percentage of CD28^+^ CD4 T cells within lung-cancer specific CD4 T cells. Objective responders (OR) are shown in green. In red, patients with no OR. Relevant statistical comparisons are shown within the graphs. Ns, no significant differences (P>0.05).

### Systemic CD8 immunity recovers in G1 responder patients following immunotherapy

In contrast to CD4 T_HD_ cells, the relative percentage of CD8 T_HD_ cells within the CD8 population did not significantly differ from age-matched healthy donors, nor could be used to identify objective responders (Supplementary Fig. 5A and 5B). Interestingly, CD8 cells from both G1 and G2 groups obtained before the start of immunotherapies did fail to proliferate after stimulation by A549-SC3 cells (Fig. 4A). To test if anti-PD-1 therapy could recover CD8 dysfunctionality *in vivo*, the proliferative capacities of CD8 T cells from G1 and G2 patients obtained after at least 3 cycles of treatment were evaluated *ex vivo* by stimulation with A549-SC3 cells. CD8 T cells from G1 responders had recovered significant proliferative capacities, while only limited enhancements were observed in G2 patients (Fig. 4B). Similarly to CD4 cells, CD8 T cells specific for lung adenocarcinoma antigens were quantified in G1 and G2 patients, and found to be comparable (Supplementary Fig. 5C) and distributed within CD28^+^ and CD28^negative^ subsets (Supplementary Fig. 5D).

**Figure 4.**
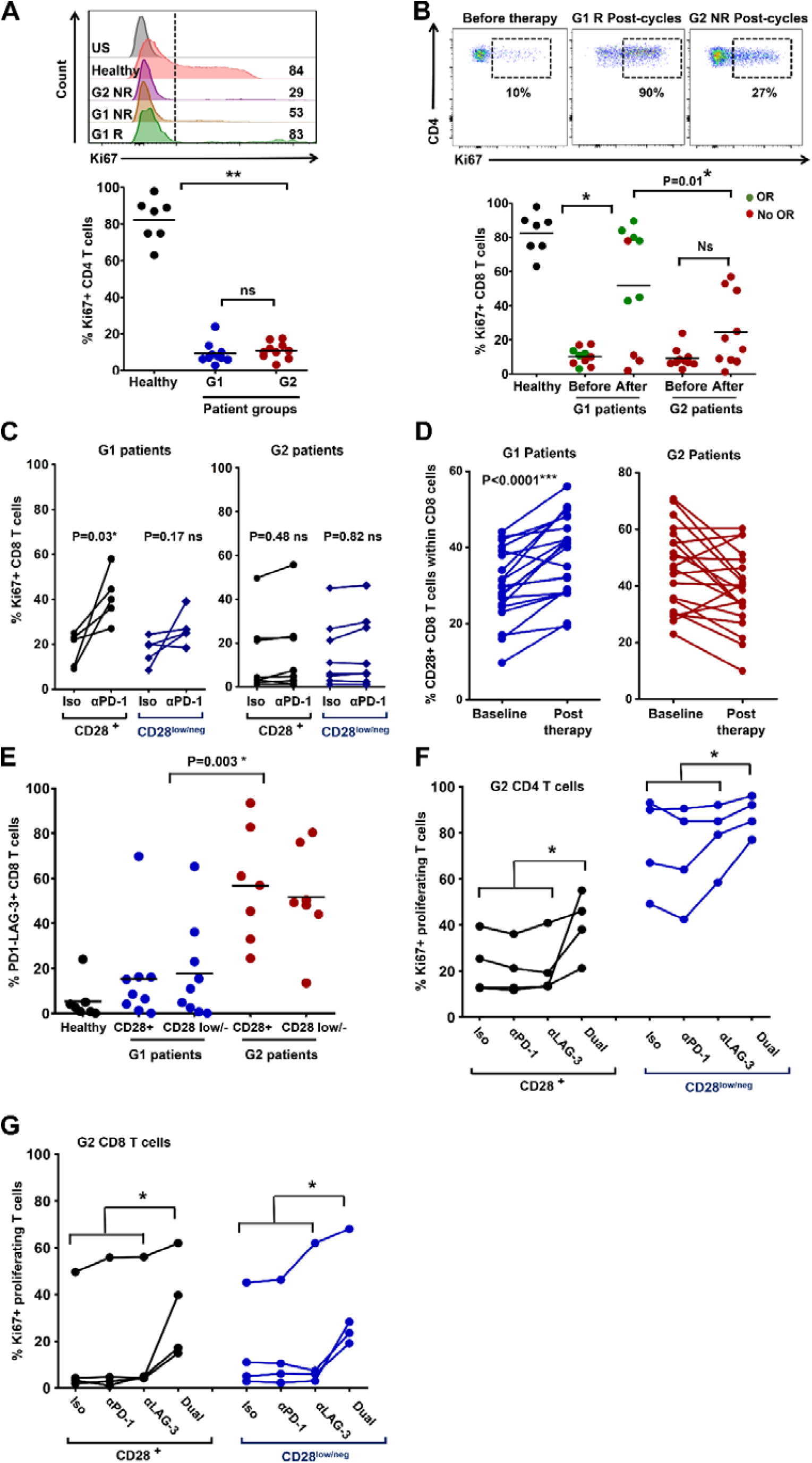
CD8 dysfunctionality in G2 NSCLC patients depends on PD-1/LAG-3 co-expression. **(A)** Expression of the proliferation marker Ki67 in CD8 T cells stimulated ex vivo by A549-SC3 cells from the indicated patient groups, before the start of immunotherapy. Relevant statistical comparisons are shown. **(B)** Dot plots of the percentage of Ki67+ proliferating CD8 T cells after ex vivo activation by A549-SC3 cells. CD8 T cells were obtained from samples of G1 or G2 patients before immunotherapy and after three cycles of anti-PD-1 therapy. Relevant statistical comparisons are shown. Green, objective responders (OR) and red, no ORs. **(C)** Same as in (A) but in the presence of an isotype control antibody or an anti-PD-1 antibody molecularly equivalent to pembrolizumab. Relevant statistical comparisons are shown. **(D)** Change in CD8 CD28^+^ T cells from baseline to post-therapy in G1 patients (left) or in G2 patients (right). **(E)** Scatter plots of PD-1/LAG-3-expressing CD8 T cells after activation by A459-SC3 cells in a sample of G1 and G2 patients within CD28^+^ and CD28^negative^ populations as indicated in the figure. **(F)** Dot-plot representing the percentage of proliferating CD4 T cells from a sample of G2 patients before starting immunotherapy, activated ex vivo by A549-SC3 cells in the presence of the indicated antibodies. “Dual” represents the addition of both anti-PD-1 and anti-LAG-3 antibodies. Appropriate statistical comparisons are shown within the graph. Data from CD28^+^ and CD28^negative^ subsets are represented as indicated. **(G)** As in (G) but for CD8 T cells.*, ** indicate significant and very significant differences, respectively. Ns, indicates non-significant differences (P>0.05).

To find out if CD8 T cells in G1 patients were especially susceptible to PD-1 blockade *ex vivo*, baseline samples of CD8 T cells from G1 and G2 patients were activated with A549-SC3 cells in the presence of an anti-PD-1 antibody or an isotype control. In agreement with the *in vivo* results, *ex vivo* PD-1 blockade improved significantly the proliferation of CD8 T cells from G1 patients, and specially CD28^+^ subsets (Fig. 4C). *In vivo* expansion of CD28^+^ CD8 T cells in murine models correlate with anti-PD-1 efficacy (12). To confirm this observation in our cohort of patients, the changes in the relative abundance of CD8 CD28^+^ T cells were compared in G1 and G2 patients from baseline to post-anti-PD-1 therapy (Fig. 4D). Accordingly, the CD28^+^ CD8 T cell compartment significantly expanded (P<0.001) only in G1 patients.

As we found that CD4 dysfunctionality correlated with high PD-1/LAG-3 co-expression in G2 patients, we tested if this was also the case for CD8 T cells. PD-1/LAG-3 co-expression was tested *ex vivo* after stimulation with A549-SC3 cells, and G2 patients presented a significantly higher proportion of PD-1/LAG-3 co-expressing CD8 T cells compared to G1 counterparts (Fig. 4E).

### CD4 and CD8 T cells from G2 patients recover proliferative activities after PD-1/LAG-3 dual blockade

Overall, our data indicated that CD4 and CD8 dysfunctionality in G2 patients was probably caused by co-expression of PD-1/LAG-3. To test if this was the case, a sample of CD4 and CD8 T cells from G2 patients before starting immunotherapies were co-incubated *ex vivo* with A549-SC3 cells in the presence of an isotype antibody control, anti-PD-1, anti-LAG-3 or anti-PD1/anti-LAG-3 antibodies. Only co-blockade of PD-1 and LAG-3 in both CD4 (Figure 4F) and CD8 T cells (Figure 4G) significantly increased the percentage of proliferating cells independently of their CD28 expression. These results confirmed that PD-1/LAG-3 co-expression strongly contributed to keeping systemic CD4 and CD8 T cells from G2 patients in a highly dysfunctional state, and that this systemic dysfunctionality can be reverted by co-blockade of both immune checkpoints.

### Objective responders are found within G1 patients with PD-L1 positive tumors

Although G1 patients exhibited functional systemic CD4 responses and TAA-specific CD4 and CD8 T cells, response rates accounted to about 50%. Hence, our results indicated that functional systemic CD4 responses were necessary but not sufficient for efficacy. Therefore, we studied other factors that contribute to responses. As NSCLC patients with high PD-L1 tumor expression benefit from anti-PD-L1/PD-1 blockade therapies (24), we assessed PD-L1 tumor expression and its association to responses in G1/G2 patients for whom PD-L1 tumor expression could be determined. Response rates of 70% were observed for G1 patients with PD-L1 positive tumors (>5%) (P=0.007) (Fig. 5A). The same benefit was observed when the stratification was extended to include G1/G2 patients with unknown PD-L1 tumor status in our cohort (Fig. 5B). PFS demonstrated a hazard ratio that favored G1/PD-L1-positivity profiling over the remaining patients in our cohort study (HR 0.187, CI95% 0.063 to 0.558; P=0.0001) with a 12-week PFS of 73.3% in contrast to 18.5% for remaining patients (P = 0.001) (Fig. 5). Remaining patients also included those for whom PD-L1 tumor expression status was unknown.

**Figure 5.**
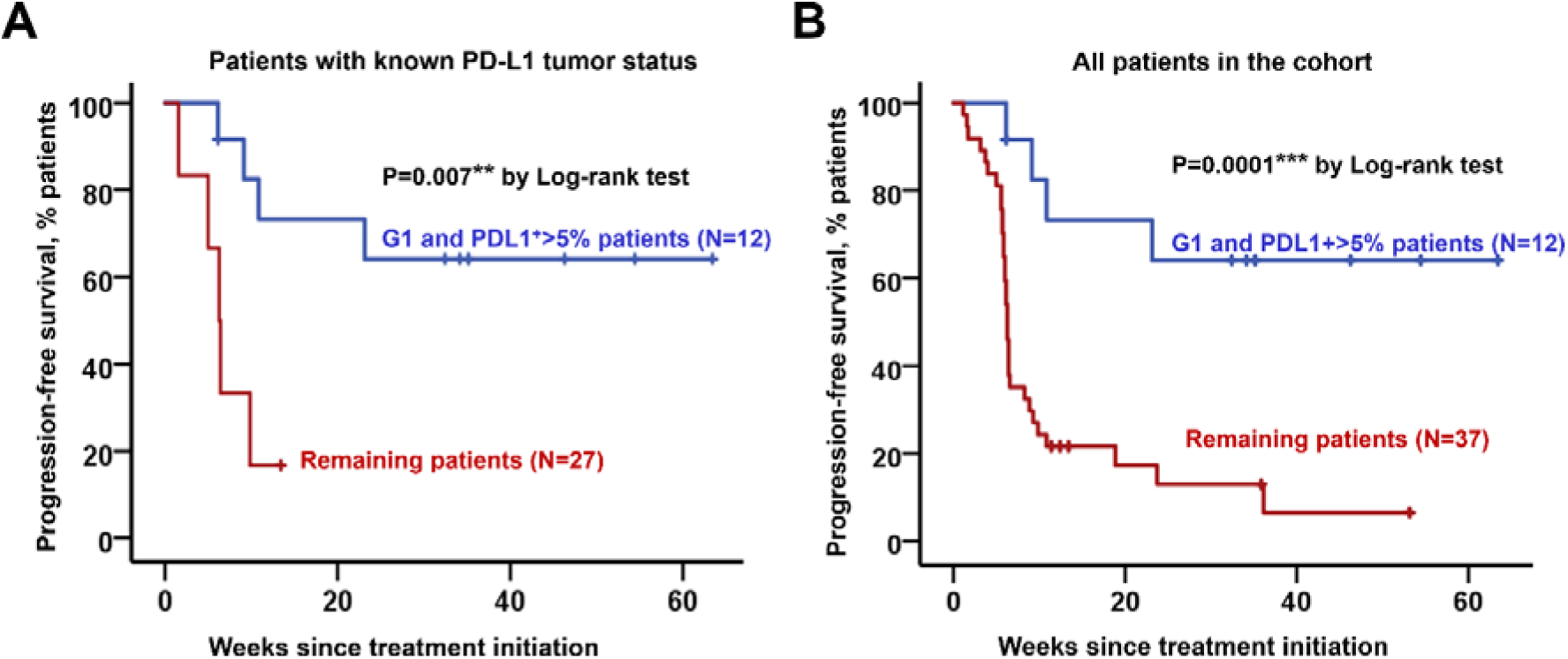
Objective responders are found within G1 patients with PD-L1+ tumors. **(A)** Kaplan-Meier plot for PFS in patients undergoing immune checkpoint inhibitor therapies stratified by G1 / PD-L1^+^ tumors (blue) and remaining patients. Only patients for whom their PD-L1 tumor status is known (red). **(B)** Same as (A) but including all patients in the study cohort. Remaining patients (red) also included G1 patients with PD-L1 low/negative tumors, G1 patients with unknown PD-L1 tumor status and G2 patients with either PD-L1^+^ or PD-L1 negative tumors.

## DISCUSSION

Tumor intrinsic and extrinsic factors contribute to the efficacy of PD-1/PD-L1 blockade therapies. So far, not a single factor has been associated to objective responses or progression, suggesting that multiple tumor-intrinsic and extrinsic mechanisms influence clinical responses. Because PD-L1/PD-1 blocking antibodies are systemically administered, these therapies cause significant and measurable systemic changes in immune cell populations (9,10). There is evidence that some of these changes may reflect the efficacy of immunotherapy in patients and used for patient stratification.

Several studies have been performed to monitor systemic dynamics of immune cell populations, some of them retrospectively and by high-throughput techniques (10–12). We evaluated responses from fresh blood samples because freezing PBMCs led to a significant alteration in the distribution of immune cell types, and distorted expression patterns of cell surface markers. Hence, sample manipulation had a significant impact on our results, which limited our study to prospective data.

Here we found that the functionality of systemic CD4 immunity is required for clinical responses to PD-L1/PD-1 blockade therapy. Indeed, it was a differential baseline factor in our cohort of NSCLC patients progressing from conventional therapies. Hence, patients with non-dysfunctional CD4 responses contained all objective responders with a response rate of about 50% (G1 patients), while no objective responses were observed in patients with highly dysfunctional CD4 T cells (G2 patients). CD4 T cell dysfunctionality in G2 patients was reflected as strongly impaired proliferation after stimulation, high co-expression of LAG3/PD-1 and resistance to *ex vivo* and *in vivo* PD-1 monoblockade. As both responders and non-responders contained comparable proportions of lung cancer-specific CD4 and CD8 T cells in our cohort of patients before the start of therapy, the experimental evidence pointed to the baseline intrinsic functionality of CD4 immunity as the key factor in our study. Importantly, patients with functional CD4 immunity could be easily identified by having a high proportion of circulating CD4 T_HD_ memory cells. ROC analysis provided a cut-off value of >40% CD4 T_hd_ to identify objective responders. We are well aware that quantification of CD4 T_HD_ cells could be used as a baseline factor for clinical stratification, pending proper independent validation which is currently ongoing. In fact, G1 patients with PD-L1 positive tumors exhibited response rates of 70%. However, the main goal of the current study was to understand the contribution of systemic T cell immunity to PD-L1/PD-1 blockade therapies, rather than providing a predictive biomarker.

Immune checkpoint inhibitor therapy aims to recover CD8 cytotoxic responses (25). To our surprise, all systemic CD8 T cells in patients before the start of immunotherapies were strongly dysfunctional. This dysfunctionality was evident by lack of proliferation following *ex vivo* activation. Nevertheless, the proliferative capacities of CD8 T cells were recovered during immunotherapy but only in patients with functional CD4 immunity. This was reflected by an expansion of CD28^+^ cells in agreement with data in murine models (12). CD8 dysfunctionality in G2 patients was again correlated with PD-1/LAG-3 co-expression. Both CD4 and CD8 dysfunctionality in G2 patients could be reverted *ex vivo* by PD-1/LAG-3 co-blockade.

An increasing number of studies are linking PD-1/LAG-3 co-expression in T cells to resistance to anti-PD-L1/PD-1 therapies (26–29). Our study suggests that patients with dysfunctional CD4 immunity should undergo alternative treatments to recover CD4 immunity and enhance CD8 T cell functionality. Our results indicate that PD-1/LAG-3 dual blockade strategies could work for G2 patients.

## METHODS

### Study Design

The study was approved by the Ethics Committee at the Hospital Complex of Navarre, and strictly followed the principles of the Declaration of Helsinki and Good Clinical Practice guidelines. Written informed consent was obtained from each participant, and samples were collected by the Blood and Tissue Bank of Navarre, Health Department of Navarre, Spain. 39 patients diagnosed with non-squamous and 12 with squamous NSCLC were recruited at the Hospital Complex of Navarre (Supplementary table 1). Patients had all progressed to first line chemotherapy or concurrent chemo-radiotherapy. Eligible patients were 18 years of age or older who agreed to receive immunotherapy targeting PD-1/PD-L1 following the current indications (Supplementary table 1). Tumor PD-L1 expression could be quantified in 39 of these patients before the start of therapies. Measurable disease was not required. The exclusion criteria consisted on concomitant administration of chemotherapy or previous immunotherapy treatment. NSCLC patients had an age of 65±8.9 (N=51). Age-matched healthy donors were recruited from whom written informed consent was also obtained, with an age of 68.60 ± 8 (mean ± S.D., N=40).

Therapy with nivolumab, pembrolizumab and atezolizumab was provided following current indications (30–32). 4 ml peripheral blood samples were obtained prior and during immunotherapy before administration of each cycle. PBMCs were isolated as described (22) and T cells analysed by flow cytometry. The participation of each patient concluded when a radiological test confirmed response or progression, with the withdrawal of consent or after death of the patient. Tumor responses were evaluated according to RECIST 1.1 (33) and Immune-Related Response Criteria (34). Objective responses were confirmed by at least one sequential tumor assessment.

### Flow cytometry

Surface and intracellular flow cytometry analyses were performed as described (3,6). T cells were immediately isolated and stained. 4 ml blood samples were collected from each patient, and PBMCs isolated by FICOL gradients right after the blood extraction. PBMCs were washed and cells immediately stained with the indicated antibodies in a final volume of 50 μl for 10 min in ice. Cells were washed twice, resuspended in 100 μl of PBS and analyzed immediately. The following fluorochrome-conjugated antibodies were used at the indicated dilutions: CD4-FITC (clone M-T466, reference 130-080-501, Miltenyi Biotec), CD4-APC-Vio770 (clone M-T466, reference 130-100-455, Miltenyi Biotec), CD4-PECy7 (clone SK3, reference 4129769, BD Biosciences,) CD27-APC (clone M-T271, reference 130-097-922, Miltenyi Biotec), CD27-PE (clone M-T271, reference 130-093-185, Miltenyi Biotec), CD45RA-FITC (reference 130-098-183, Miltenyi Biotec), CD62L-APC (reference 130-099-252, Miltenyi Biotech), CD28-PECy7 (clone CD28.2, reference 302926, Biolegend), PD-1-PE (clone EH12.2H7, reference 339905, Biolegend), CD8-FITC (clone SDK1, reference 344703, Biolegend), CD8-APC-Cy7(clone RFT-8, reference A15448, Molecular probes by Life technologies), CD57-PE (clone HCD57), reference 322311, Biolegend), H2AX-FITC (clone 2F3, reference 613403, Biolegend), IL-2 Alexa Fluor 647 (clone MQ1-17H12, reference 500315, Biolegend), IFN δ (clone 4S.B3), reference 50256, Biolegend). **CD14-brilliant blue. CD3-**

### Cell culture

Human lung adenocarcinoma A549 cells were a kind gift of Dr Ruben Pio, and were grown in standard conditions. These cells were modified with a lentivector encoding a single-chain version of a membrane bound anti-OKT3 antibody (35). The lentivector expressed the single-chain antibody construct under the control of the SFFV promoter and puromycin resistance from the human ubiquitin promoter in a pDUAL lentivector construct (6). The single-chain antibody construct contained the variable light and heavy OKT3 immunoglobuline sequences separated by a G-S linker fused to a human IgG1 constant region sequence followed by the PD-L1 transmembrane domain.

Monocyte-derived DCs were generated from adherent mononuclear cells in the presence of recombinant GM-CSF and IL-4 as described (22). DCs were loaded with A549 protein extract obtained after three cycles of freezing/thawing. Loading was carried out overnight, and DCs were matured with 10 ng/ml of IFN-γ before adding T cells in a 1:3 ratio as described (22).

### Anti-PD-1 antibody production and purification

To generate an antibody molecularly equivalent to the published sequence of pembrolizumab, a cDNA encoding the published aminoacid sequences of the heavy and light immunoglobulin chains (21) were cloned and expressed in chinese hamster ovary (CHO) cells. Supernatants were collected and antibodies purified by affinity chromatography following standard procedures.

### Data collection and Statistics

T cell percentages were quantified using Flowjo (17,18). The percentage of CD4/CD8 T_HD_ (CD28 and CD27 double-negative) and poorly differentiated T cells (CD28+ CD27+) were quantified prior to therapy (baseline), and before administration of each cycle of therapy within CD4 and CD8 cells. Gates in flow cytometry density plots were established taking non-senescent T cells as a reference. Data was recorded by M.Z.I., and separately analyzed thrice by M.Z.I. and H.A.E. independently. Cohen’s kappa coefficient was utilized to test the inter-rater agreement in classification of immunological profiles (κ =0.939).

The mode of action, pharmacokinetics, adverse events and efficacies of the three PD-L1/PD-1 blocking agents are comparable in NSCLC, which act through the interference with the inhibitory interaction between PD-L1 and PD-1 (30–32). Treatments administered to the patients were allocated strictly on the basis of their current indications, and independently of any variable under study. All data was pre-specified to be pooled to enhance statistical power, and thereby reducing type I errors from testing the hypotheses after *ad hoc* subgrouping into specific PD-L1/PD-1 blockers. The number of patients assured statistical power for Fisher’s exact test of 0.95 and superior for Student t and Mann-Whitney tests (G*Power calculator) (36), taking into account that the expected proportion of responders is around 25% to 35% (if no stratification using PD-L1 tumor expression levels is used) (30–32). Two pre-specified subgroup analyses in the study were contemplated. The first, baseline T cell values; the second, post-first cycle T cell changes from baseline. The study protocol contemplated the correlation of these values with responses using Fisher’s exact test, paired Student t tests/repeated measures ANOVA (if normally distributed) or U of Mann-Whitney/Kruskal-Wallis (if not normally distributed, or data with intrinsic high variability) to be correlated with responses. Two-tailed tests were applied with the indicated exceptions (see below). Importantly, the treatment administered to the patients was allocated independently of their basal immunological profiles, and strictly followed the current indications for the PD-L1/PD-1 inhibitors.

The percentage of T cell subsets in untreated cancer patients was normally distributed (Kolmogorov-Smirnov normality test), but not in age-matched healthy donors. Hence, to compare T cell values between two independent cancer patient groups, two-tailed unpaired Student t tests were used, while comparisons between healthy subjects and cancer patients were carried out with the U of Mann-Whitney. Percentages of T cell populations in treated patients were not normally distributed, so response groups were compared with either Mann-Whitney (comparisons between two independent groups) or Kruskall-Wallis for multi-comparison tests if required. To test changes in Ki67 expression in T cells between control and treatment paired groups, two-tailed paired t tests were carried out. Fisher’s exact test was used to assess the association of the baseline values of T_HD_ cells with clinical responses. The same tests were performed to assess associations between G1/G2 groups and neutrophil-to-lymphocyte ratios and also the indicated patient characteristics. *Post hoc* Cohen’s kappa coefficient test was used to test the agreement of radiological versus immunological criteria for the identification of hyperprogressors.

Progression free survival (PFS) was defined as the time from the starting date of therapy to the date of disease progression or the date of death by any cause, whichever occurred first. PFS was censored on the date of the last tumor assessment demonstrating absence of progressive disease in progression-free and alive patients. PFS rates at 12 and 28-weeks was estimated as the proportion of patients who were free-of-disease progression and alive at 12 and 28 weeks after the initiation of immunotherapies. Patients who dropped out for worsening of disease and did not have a 28-week tumor assessment were considered as having progressive disease. Overall response rate (ORR) was the proportion of patients who achieved best overall response of complete or partial responses.

PFS was represented by Kaplan-Meier plots and long-rank tests utilized to compare cohorts. Hazard ratios were estimated by Cox regression models. For PFS analyses, patients were prespecified to be stratified only on the basis of their basal T_HD_ values to avoid increase in type I errors due to multiplicity by subgroup analyses. Receiver operating characteristic (ROC) analysis was performed with baseline T_HD_ numbers and response/no response as a binary output. Statistical tests were performed with GraphPad Prism 5 and SPSS statistical packages.

### Validation dataset

Data from a set of 32 patients was validated in parallel by independent handling, processing, staining, flow cytometry data collection and analysis. The validation dataset was generated by a technician working in unrelated research themes. For the validation dataset, a previous step of monocyte depletion by adherence was performed in some to the samples to eliminate background from CD14^+^ CD4^low^ monocytic cells. CD14^+^ cells were gated out for the rest of the samples. *Post hoc* Cohen’s kappa coefficient test was used to test the agreement between the discovery cohort *versus* the validation cohort on classification of G1/G2 patients.

## Acknowledgments

We sincerely thank the patients and families that generously agreed to take part in this study. We are thankful to Drs Luis Montuenga and Ruben Pio for their constructive comments and input.

## Author contributions

M.Z.I. designed and carried out experiments, collected data, analyzed data. H.A. designed and carried out experiments, collected data, analyzed data. G.F. recruited patients, collected data, analyzed clinical data. MJ G. C., M.G.C. and A.B. carried out experiments, collected data, analyzed data. M.M, B.H., L.T., I.M., M.J.L., A.F.L. and R.V. recruited patients, collected data, analyzed clinical data. M.J.L. recruited patients, collected data, analyzed clinical data. A.F. recruited patients, collected data, analyzed clinical data. L.T. recruited patients, collected data, analyzed clinical data. R.V. supervised the clinical staff, recruited patients, analyzed clinical data. G.K. conceived the project, supervised non-clinical researchers, analysed data and wrote the paper. D.E. conceived the project, supervised non-clinical researchers, analysed data and wrote the paper. All authors participated in the writing of the manuscript.

## SUPPLEMENTARY TABLES

**Supplementary table 1.**
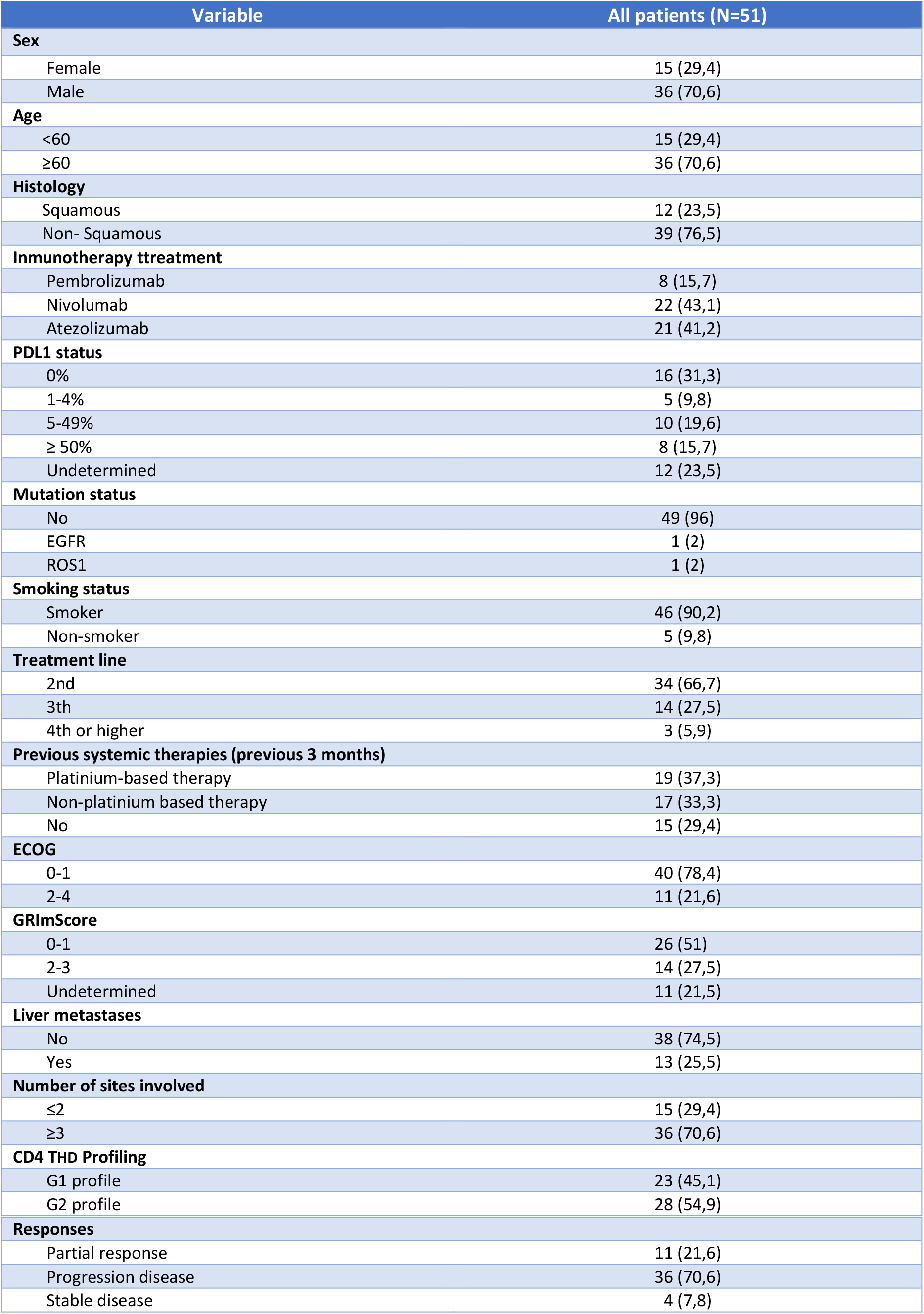
Baseline patient characteristics.

**Supplementary table 2.**
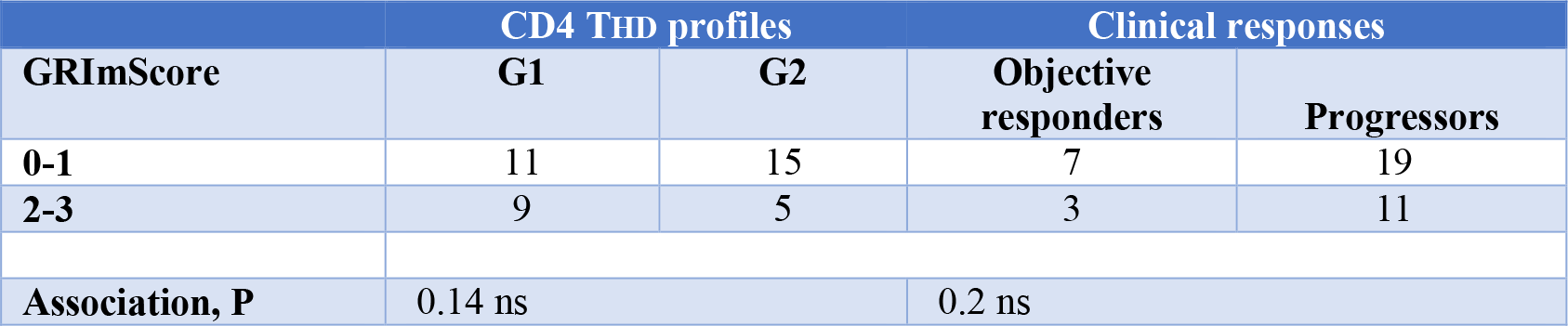
Association of CD4 T cell profiles with GRIm score.

## SUPPLEMENTARY FIGURES

**Supplementary figure 1.**
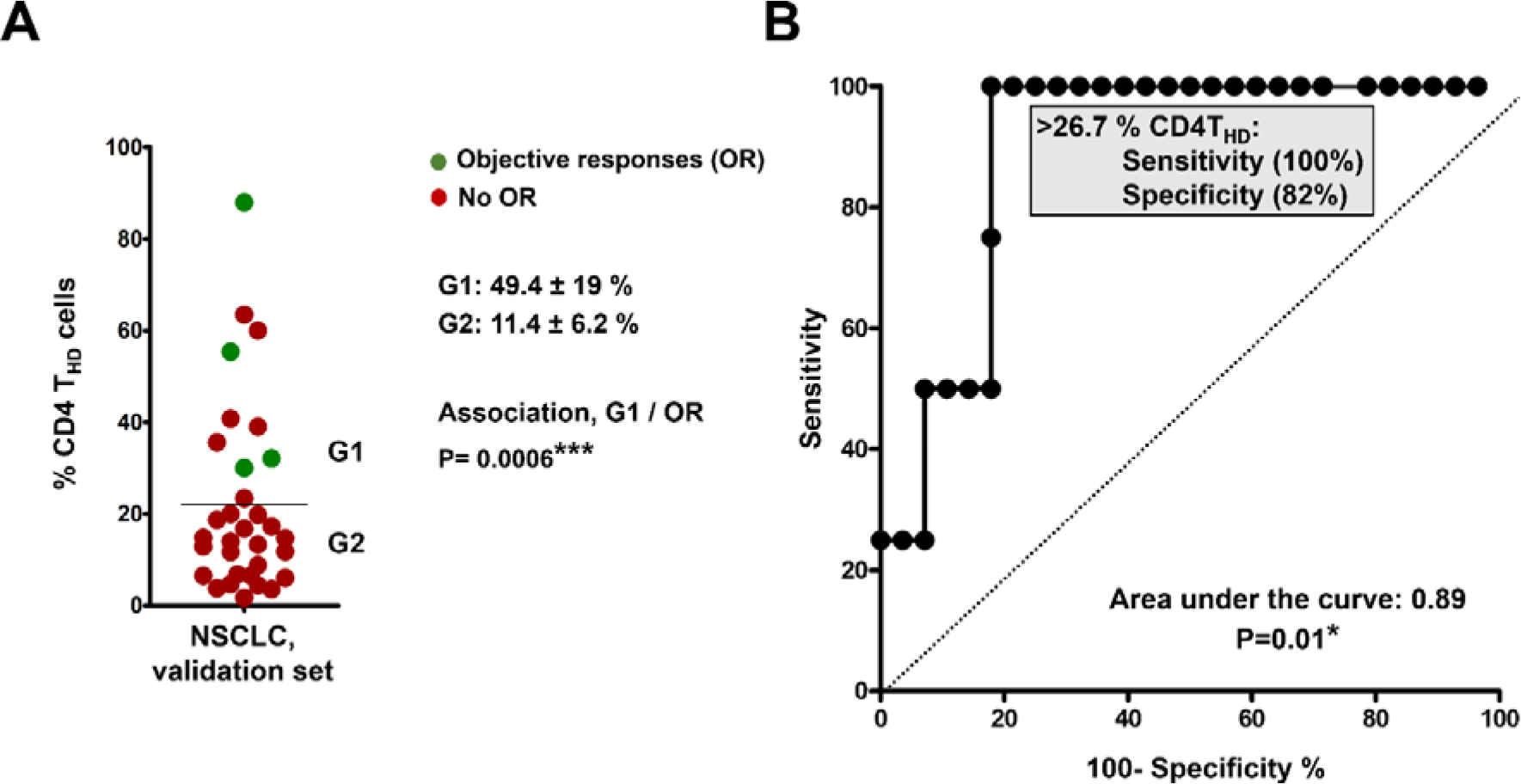
Validation dataset. **(A)** Distribution of circulating CD4 T_HD_ cells within CD4^+^ CD14^negative^ cells in NSCLC patients constituting the validation set (N=32). G1 and G2 groups are indicated and separated by the mean (horizontal line). The means ± standard deviations of CD4 T_HD_ cells in G1 and G2 groups are shown on the right, as well as the association between G1 profiles and objective responses by the Fisher’s test. **(B)** ROC analysis of CD4 THD quantification in the validation dataset and objective responses. The cut-off value for identification of responses is shown in the graph. *, **, indicate significant (P<0.05) and highly significant (P<0.001) differences.

**Supplementary Figure 2.**
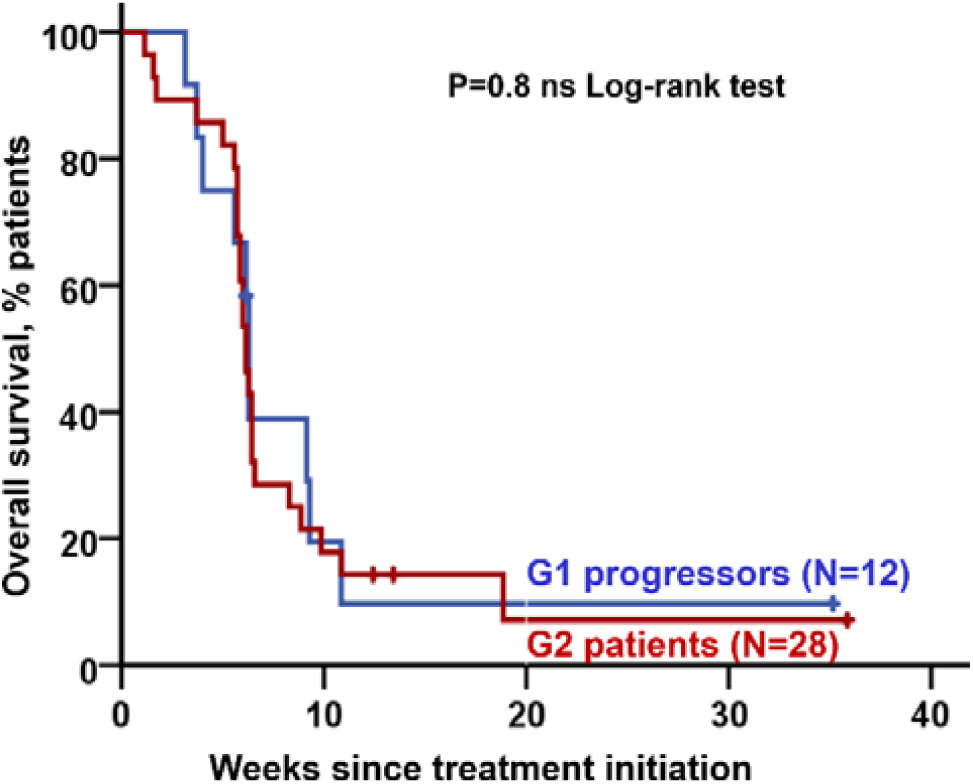
CD4 T cell profiling does not have significant prognostic value. Kaplan-Meier plot of overall survival (OS) for G1 progressors (blue) and G2 patients (red), as indicated.

**Supplementary figure 3.**
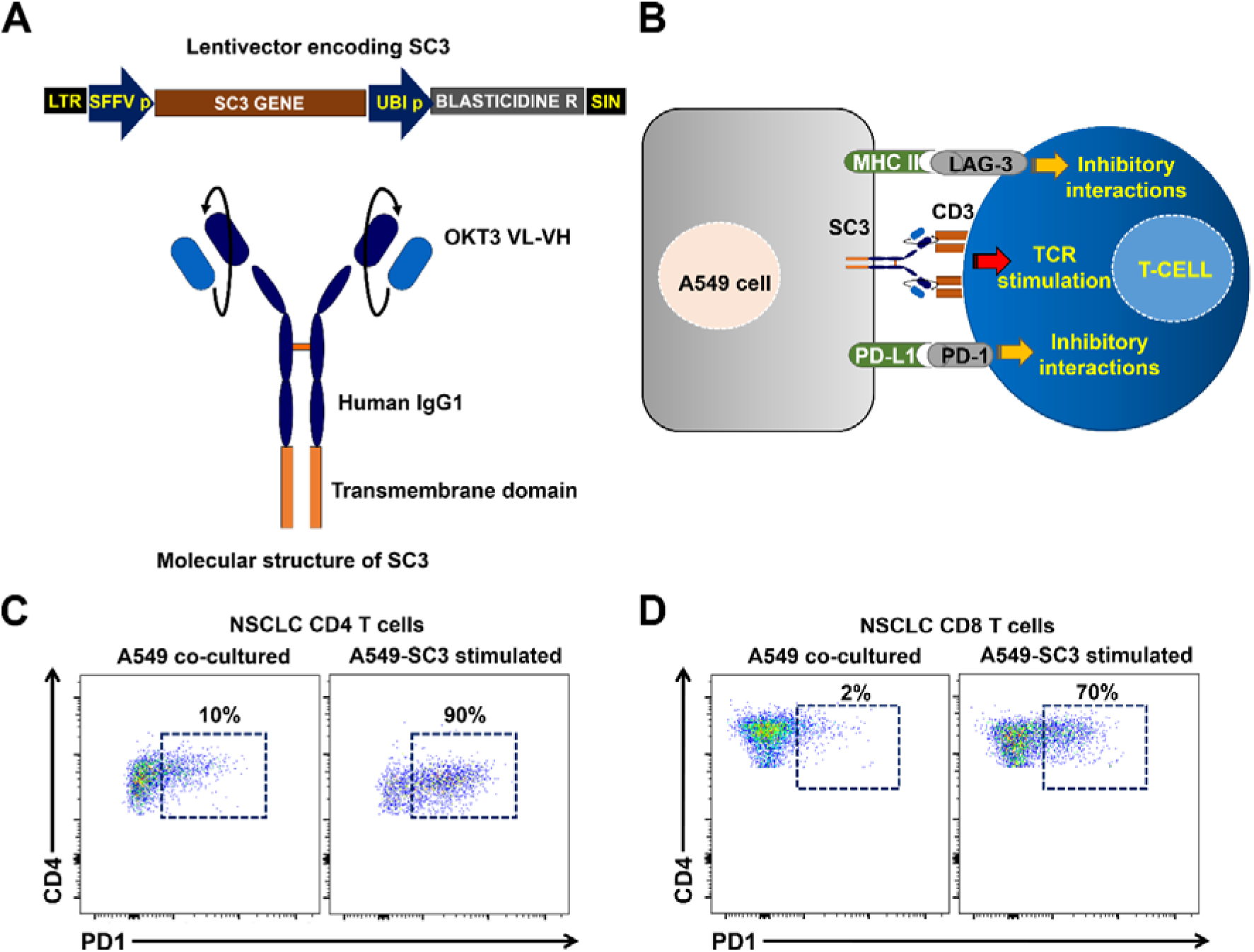
Ex vivo human lung adenocarcinoma-T cell recognition system. **(A)** Top, lentivector co-expressing an anti-CD3 single chain antibody gene (SC3) and blasticidine resistance for selection. SFFVp, spleen focus-forming virus promoter; UBIp, human ubiquitin promoter; LTR, long terminal repeat; SIN, U3-deleted LTR leading to a self-inactivating lentivector. Bottom, molecular structure of the SC3 molecule, which is anchored to the cell membrane by a transmembrane domain as indicated. OKT3 VL, variable region of the light chain from the anti-CD3 antibody OKT3; VH, variable region of the heavy chain from the anti-CD3 antibody OKT3. **(B)** Scheme of the cell-to-cell interactions mediated by the lentivector-modified A549 cell and T cells including SC3/CD3, PD-L/PD-1 and MHCII/LAG-3 interactions as indicated. **(C)** Representative flow cytometry density plots with the up-regulation of PD-1 expression in CD4 from NSCLC patients following co-incubation with A549-SC3 cell as indicated (right graph), or with unmodified A549 control (left graph). Percentages of PD-1+ T cells are shown within the graphs. **(D)** As in (C) but with CD8 T cells. Percentages of PD-1+ T cells are shown within the graphs.

**Supplementary Figure 4.**
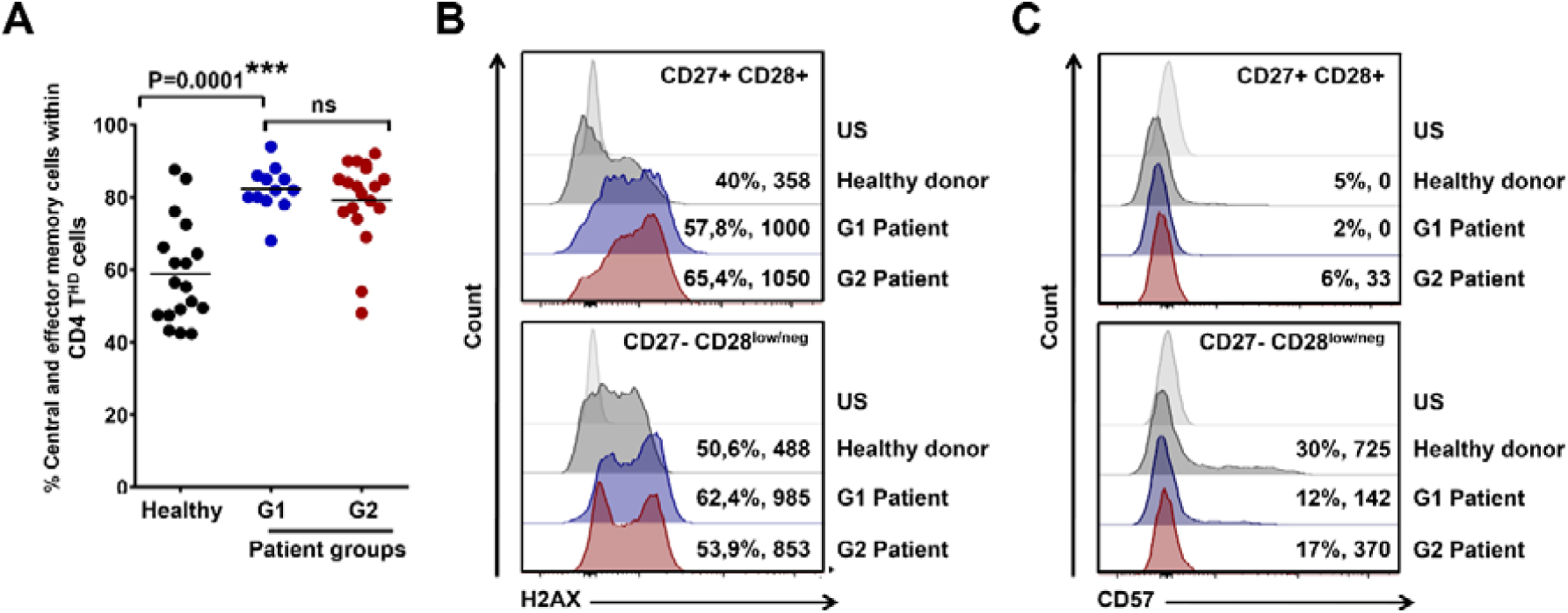
CD4 THD cells in NSCLC patients are mainly non-senescent memory subsets. (A) Scatter plot graphs of the percentage of memory phenotypes in baseline CD4 T_HD_ cells according to CD62L-CD45RA expression (% CD45RA^negative^ CD62L^positive^ central memory + % CD45RA^negative^ CD62L^negative^ effector memory cells) in a sample of healthy donors, G1 and G2 patients, and in healthy donors as indicated. **(B)** Expression of the genotoxic damage maker H2AX by flow cytometry in CD4 T cell subsets from an aged-matched healthy donor, and NSCLC G1 and G2 patients as indicated. Percentage of positivity and mean fluorescent intensities are indicated for each population. Top, histogram analysis within CD27+ CD28+ CD4 T cells, and bottom, CD27^negative^ CD28^low/negative^ counterparts as indicated. **(C)** As in (B) but for CD57 expression. US, unstained control; NS, no significant differences (P>0.05); ***, highly significant differences (P<0.001).

**Supplementary Figure 5.**
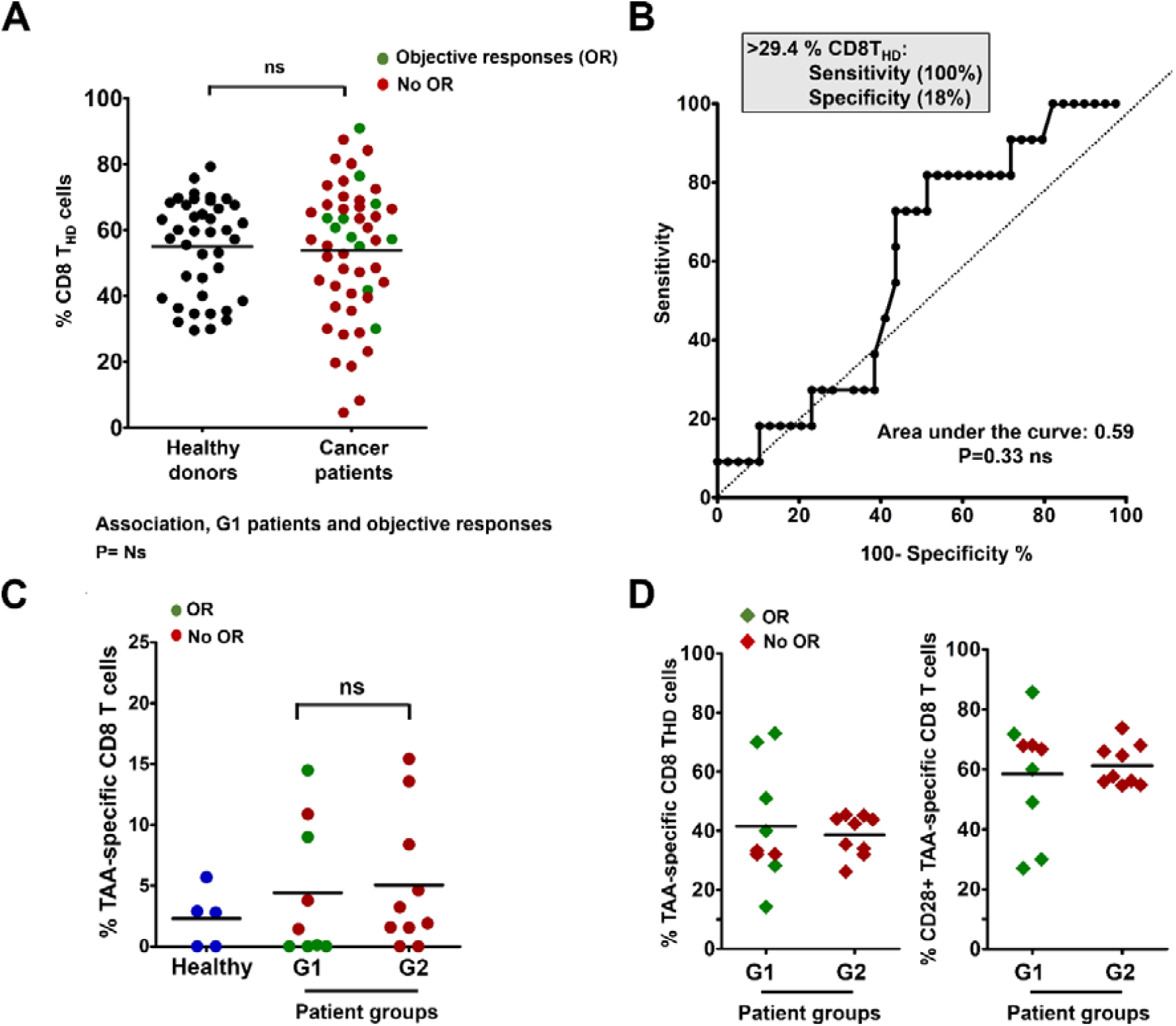
Systemic CD8 responses in NSCLC patients. **(A)** Percentage of circulating highly differentiated CD8 cells in age-matched healthy donors and NSCLC patients before undergoing immunotherapies. Relevant statistical comparisons are shown. In green, objective responders (OR). In red, no OR. **(B)** ROC analysis of baseline CD8 T_HD_ quantification as a function of objective clinical responses. **(C)** Dot-plot of lung cancer antigen-specific CD8 T cells obtained before the start of immunotherapies and stimulated with A549-loaded autologous DCs in G1 and G2 patients, as indicated. **(D)** Left dot plot, percentage of CD28-negative CD8 T cells within TAA-specific CD8 subsets in G1 and G2 patients as indicated. Right dot plot, same as left but with CD28-positive subsets. Green, objective responders (OR) and red, no ORs.

